# Mechanisms of a sustained anti-inflammatory drug response in alveolar macrophages unraveled with mathematical modeling

**DOI:** 10.1101/2020.04.13.031245

**Authors:** Elin Nyman, Maria Lindh, William Lövfors, Christian Simonsson, Alexander Persson, Daniel Eklund, Erica Bäckström, Markus Fridén, Gunnar Cedersund

**Affiliations:** Department of Biomedical Engineering, Linköping University, Linköping, Sweden; Drug Metabolism and Pharmacokinetics, Research and Early Development, Respiratory, Inflammation and Autoimmune (RIA), BioPharmaceuticals R&D, AstraZeneca, Gothenburg, Sweden; Inflammatory Response and Infection Susceptibility Centre (iRiSC), Department of Medical Sciences, School of Medicine and Health, Örebro University, SE-701 82 Örebro, Sweden; Translational PKPD Group, Department of Pharmaceutical Biosciences, Uppsala University, Uppsala, Sweden

## Abstract

Both initiation and suppression of inflammation are hallmarks of the immune response. If not balanced, the inflammation may cause extensive tissue damage, which is associated with common diseases, e.g. asthma and atherosclerosis. Anti-inflammatory drugs often come with severe side effects driven by high and fluctuating drug concentrations. To remedy this, drugs with sustained anti-inflammatory responses are desirable. However, we still lack a quantitative mechanistic understanding of such sustained effects. Here, we study the anti-inflammatory response to a common glucocorticoid drug, Dexamethasone. We find a sustained response 22 hours after drug removal. With hypothesis testing using mathematical modeling, we unravel the underlying mechanism – a slow release of Dexamethasone from the receptor-drug complex. The developed model is in agreement with time-resolved training and testing data, and is used to simulate hypothetical treatment schemes. This work opens up for a more knowledge-driven drug development, to find sustained anti-inflammatory responses and fewer side effects.

## Introduction

The inflammatory response against infections rely on activation of the innate immune system. This activation contributes to a temporal induction of cytokines and various other specific signaling molecules, in turn attracting and instructing additional immunocompetent cells. This response is fast and rely on both local production and massive recruitment of immunocompetent cells, which are directed to the site of inflammation from the blood stream. This proinflammatory process needs to be restricted by anti-inflammatory mediators in order to return to homeostasis and to avoid extensive tissue damage caused by the inflammation [1]. When this balance act fails, common human disease states, such as septic shock, asthma, rheumatoid arthritis, inflammatory bowel diseases, multiple sclerosis, and atherosclerosis occur [2–4]. To control the proinflammatory mechanisms of such diseases, anti-inflammatory drugs that target several specific and non-specific mechanisms have been on the market for decades. However, such anti-inflammatory drugs can cause severe side effects in the gastrointestinal tract, liver and kidney, as well as allergic reactions and edemas. Therefore, drugs with a sustained response are attractive, since side effects could be minimized by lowering the fluctuation and/or the peak level of plasma drug concentration. Further, if we understand the mechanisms of such a sustained response, we can use this knowledge in the search for new and better drug candidates.

Anti-inflammatory drugs act to reduce the production of cytokines. Cytokines are small signaling molecules central for directing down-stream immunological effects. Some cytokines act on specific pathways and cell types whereas others are broader in their span of activity. Tumor necrosis factor (TNF) is a hallmark proinflammatory cytokine acting through both paracrine and autocrine pathways. TNF mediates inflammatory activation in a multitude of ways. For example, TNF is involved in causing fever, limiting viral replication, increasing phagocytic cells’ capacity to kill pathogens, as well as in stimulating cells to release more cytokines and chemokines. This increased release attracts leukocytes and other cells which in turn further propagate the inflammatory process [5]. TNF is produced by both immunocompetent cells of myeloid and lymphocyte lineage as well as non-immunocompetent cells such as keratinocytes, endothelial cells, and neurons. One of the main contributors of soluble TNF in the inflammatory setting is the macrophage [6,7]. This is especially true for alveolar macrophages, which are highly specialized macrophages present in the lungs. The alveolar macrophages function primarily in the defense against, and in the response to, inhaled particles and potentially pathogenic microorganisms. Macrophages thus play a critical role in the pathophysiology of inflammatory lung disease, such as asthma, and cystic fibrosis. In summary, one of the known pathways by which inhaled particles or microorganisms stimulate the recruitment, and subsequent activation, of inflammatory cells in lung disease, is through the activation of these alveolar macrophages to produce and release TNF.

TNF production is under strict transcriptional control where activation of the cell leads to an expansion of the TNF mRNA pool, which gives rise to rapid translation and eventual secretion of TNF, which in turn are free to act on surrounding tissues and cells (Figure 1, left). A major regulator of TNF transcription is the transcription factor: nuclear factor *κ*-light-chain-enhancer of activated B cells (NF-*κ*B). NF-*κ*B is always present in an inactive state in the cytoplasm of the cell, bound to the inhibitor of NF-*κ*B, I*κ*B (I*κ*B*α*) [8, 9]. Upon certain cellular activation, e.g. through toll like receptor (TLR) 4, I*κ*B become phosphorylated by the I*κ*B kinase (IKK)-complex leading to degradation by the proteasome. This in turn renders NF-*κ*B active and capable of translocation to the nucleus, where it promotes transcription of multiple genes with for proinflammatory function, among them TNF. In the signaling pathway, there are several more proteins involved, e.g. Interleukin-1 receptor-associated kinase (IRAK) enzymes, and TNF receptor associated factor (TRAF) proteins (see Figure 1, left). Finally, there are drugs that interfere with the NF-*κ*B activity, among them glucocorticoids.

**Figure 1:**
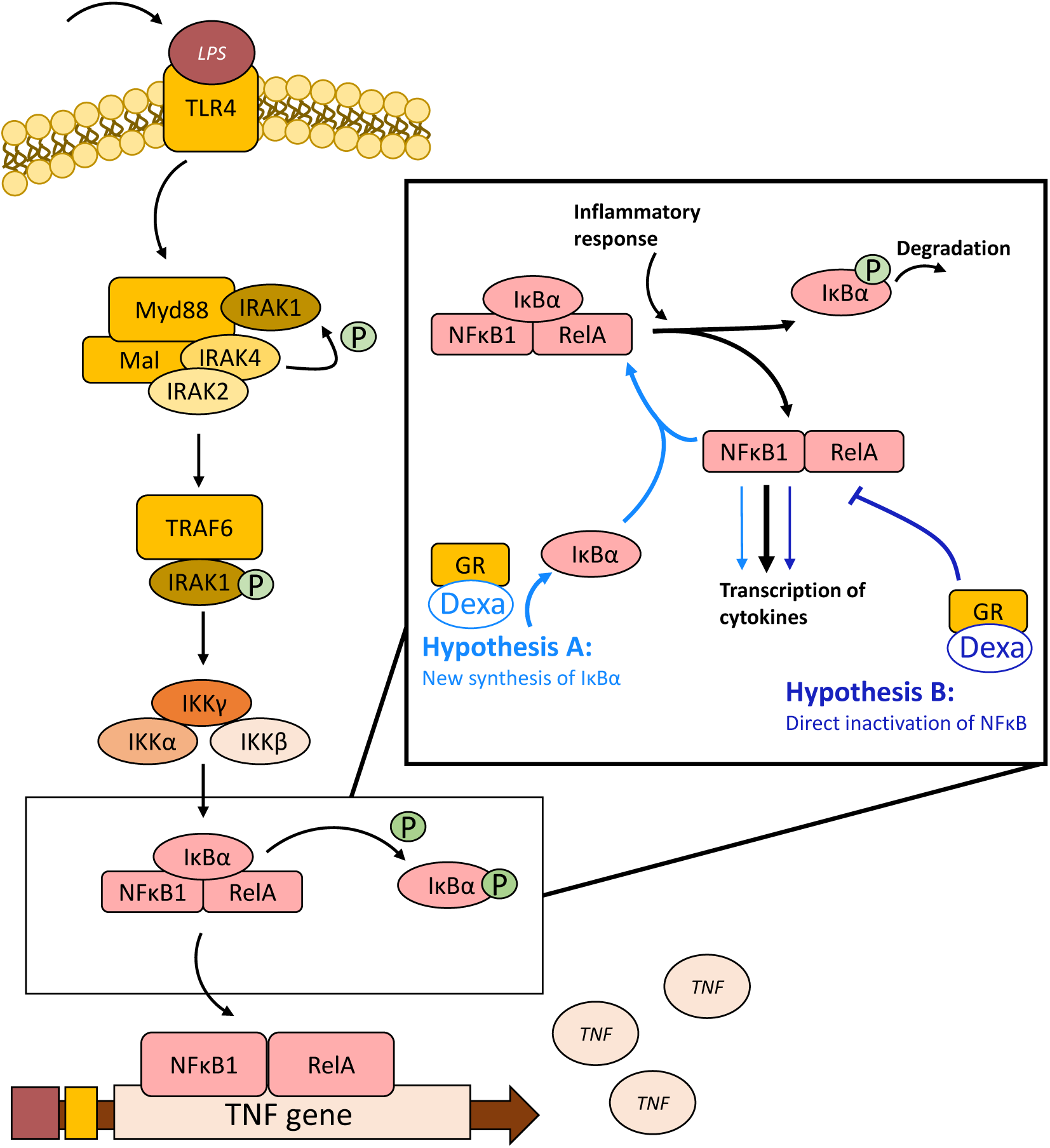
The inflammatory signaling pathway in macrophages. LPS stimulates TLR4 which leads to a cascade of signaling events, resulting in the transcription and release of TNF*α*. The squared box contains both hypotheses for the effect of the anti-inflammaroty drug Dexamethasone (Dexa). In Hypothesis A, the protein I*κ*B*α* is synthesized in response to Dexa bound to glucocorticoid receptors (GR). I*κ*B*α* in turn binds to NF-*κ*B proteins (RelA and NF-*κ*B1) to inhibit their transcriptional activity. In hypothesis B, there is a direct physical association between activated GR and the NF-*κ*B subunit RelA that reduces the transcriptional activity.

Glucocorticoids, e.g. cortisol, counteract inflammatory responses, and several synthetic glucocorticoids have been developed for treatment of inflammatory diseases. Glucocorticoids act through the glucocorticoid receptor (GR) and inhibits the release of cytokines from macrophages, e.g. through inhibition of transcription, changes in mRNA stability, changes in protein translation and/or post-translational processing. The synthetic glucocortocoid Dexamethasone (Dexa) is 30 times more potent than the endocrine glucocorticoid cortisol in inhibiting cytokine production [10]. There are two main hypotheses for the mechanisms of action of Dexa (Figure 1, box to the right): Hypothesis A: new synthesis of the protein I*κ*B that binds to NF-*κ*B and thereby hinders the inflammatory response [11,12], and Hypothesis B: a physical association between activated GR and the NF-*κ*B subunit p65/RelA that reduces the activity of NF-*κ*B [13, 14]. Hypothesis B is potentially a general mechanism for many cell types. Hypothesis A has been shown e.g. in Jurkat cells transfected with GR were I*κ*B mRNA levels were increased at 30-60 minutes after addition of Dexa [11], while this mechanism has been shown to be lacking in endothelial cells [15]. Neither of these hypotheses have been formally tested with e.g. a mathematical modeling framework.

Mathematical modeling is used in several fields of science to analyse data in a systematic and quantitative way. In the field of biology, modeling methods are commonly referred to as systems biology, and often focus on intracellular metabolic and/or protein signaling pathways. In the field of pharmacology, pharmacokinetic/pharmacodynamic (PK/PD) modeling is more commonly used, which instead often focus on drug-receptor binding and the selection of optimal drug doses. The combination of both approaches, i.e. the use of models of drug actions, with a clear biological interpretation that can be used to gain mechanistic insights, e.g. regarding intracellular signaling is commoly referred to as systems pharmacology [16].

Previous efforts to model TNF secretion from macrophages includes both systems biology and PK/PD models (see [17] for a review of models). For example, systems biology modeling has been used for a detailed elucidation of the role of different TLRs and their ligands, including bacterial lipopolysaccharides (LPS), in both endocrine and paracrine TNF signaling [18]. This model, however, does not contain the inhibitory effects of glucocorticoids and the mechanism of such an inhibition. Hao and co-workers [19] have developed a model of chronic pancreatitis to simulate the effect of disease-modifying agents. This model includes the pancreatic micro-environment, including cytokines and macrophages. This model, however, does not contain the details of the intracellular signaling pathways within macrophages, and instead the interplay between different players of the micro-environment is targeted. Mechanistic PK/PD models for glucocorticoid receptor signaling have been developed for the metabolic side-effect in the liver, mediated via tyrosine aminotransferase [20]. However, no existing model can be used to study: i) the different hypotheses of Dexa-induced anti-inflammation, and ii) potential intracellular mechanisms behind a sustained anti-inflammatory response.

Here, we study the anti-inflammatory response of alveolar macrophages to Dexa, and find a sustained cellular response to the drug up until 22 hours after withdrawal of the drug. To unravel the mechanisms behind such a sustained response, we use a mathematical modeling approach. First, we test the different hypothesis for the intracellular action of Dexa, and reject Hypothesis A: “New synthesis of I*κ*B*α*”. Hypothesis B: “Direct inactivation of NF-*κ*B”, on the other hand, is in agreement with all our time-resolved data series. The mechanism behind the sustained response in Hypothesis B is a slow release of Dexa from the GR-Dexa complex. We use the final model to simulate different treatment schemes and find that the sustained response of Dexa allows for a one-dose-daily treatment scheme, which could be beneficial to reduce side effects of a drug.

## Method

### Experimental methods

All experimental methods are found in the Supplementary Material.

### Mathematical modeling

A system of ordinary differential equations is used to model the dynamic response to LPS and Dexa in alveolar macrophages. The same model structure is used for LPS stimulated inflammation for both hypotheses:

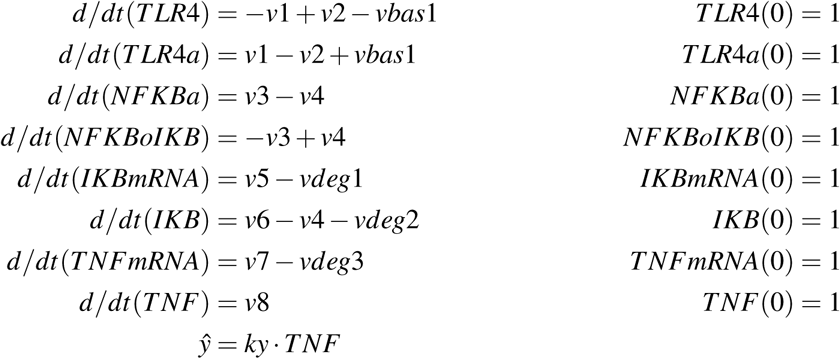

where e.g. *T LR*4 and *T LR*4*a* are states with initial conditions specified as *T LR*4(0) and *T L R*4*a*(0). *ŷ* corresponds to the measured output. *v*1, *v*2, etc are the reaction rates, further defined as:

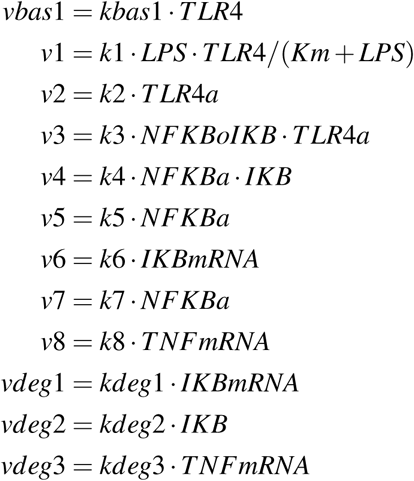

where e.g. *kbas*1 and *k*1 are parameters with unknown values.

The anti-inflammatory effect of Dexa is implemented in different ways for Hypothesis A and B. The activation of the receptor complex *DexaGR* is modeled in the same way for both hypotheses:

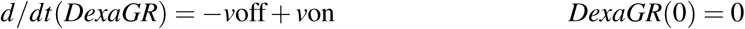

where

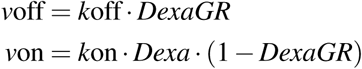

In Hypothesis A, *DexaGR* increases the rate of formation of *IKBmRNA*:

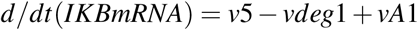

where

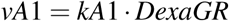

In Hypothesis B, *DexaGR* is activated in the same way as in Hypothesis A, and the effect of the complex is to inhibit *v*7:

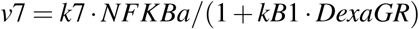

An interaction graph that depicts the models behind both Hypothesis A and B is showed in Figure 2.

**Figure 2:**
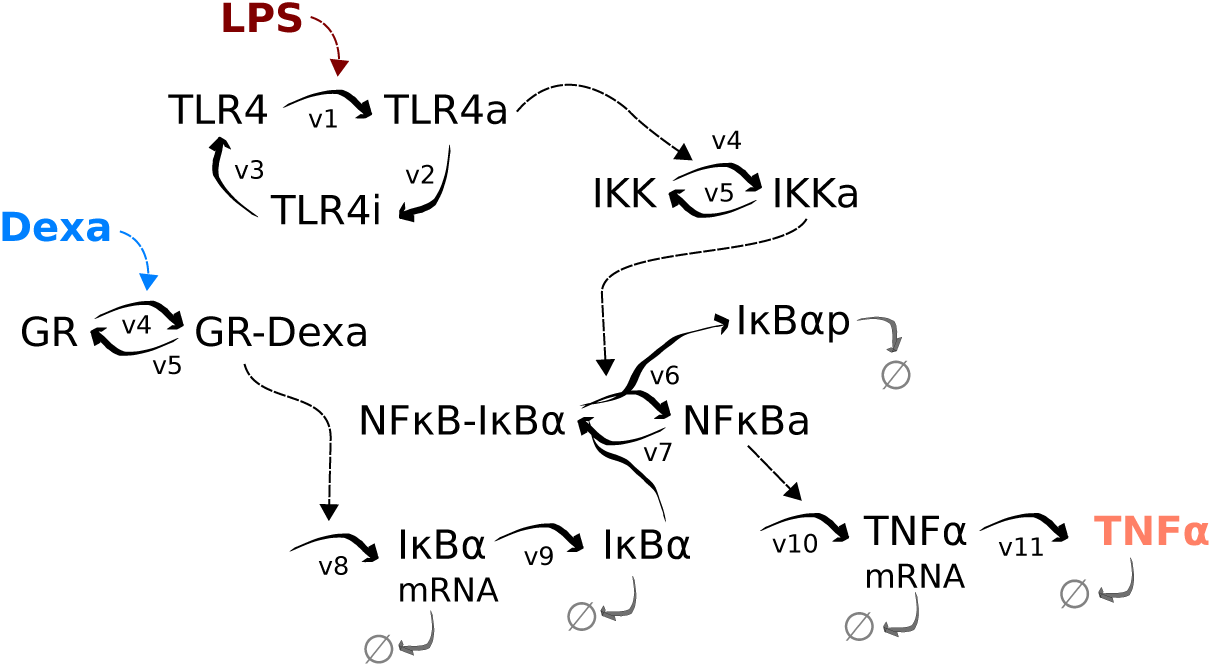
The developed mathematical models for Hypothesis A and B, both with 9 states and 16 parameters. Model inputs are lipopolysaccaride (LPS) and Dexamethasone (Dexa), model output is tumor necrosis factor (TNF). Thick arrows represent flows and thin dashed arrows represent activating signals. GR - glucocorticoid receptor, IKK - I*κ*B kinase, I*κ*B*α* - inhibitor of *κ* B, TLR4 - Toll-like receptor 4, NF-*κ*B - nuclear factor *κ*-light-chain-enhancer of activated B cells, p - phosphorylation, a - active state.

### Parameter estimation

Most rate constants were allowed a free range (1e-3, 1e3), but for *k*on and *k*off in the Dexa-GR binding there are data available that restrict the rate for *k*off to 0.001 - 0.01 /min and *k*on to 0.5 - 1 /*µ*M/min and (different amounts of GR were expressed in COS1 cells) [21]. The range give rise to a range of affinity of *Kd* = 0.001 - 0.02 *µ*M, which is consistent with competition experiments in THP-1 cells [22] and ex vivo lung tissue from rats [23]. However, another measurement for *k*off is 0.0034 - 0.007 /s, i.e. 0.21 - 0.42 /min (GR expressed in High Five cells) [24]. The non-overlapping measurements made us allow for a wide range (1e-3, 1e3) also for *k*on and *k*off, and allowed the parameter estimation to find the best possible values also for these rate parameters.

The agreement between model simulations and data is quantified with a cost function, *V* (*p*)

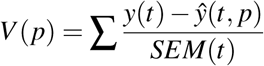

where the sum is over all measured time points, *t*; *p* is the parameters; *y*(*t*) is the measured data and *ŷ*(*t*) is the model simulations; *SEM*(*t*) is the standard error of the mean for the measured data.

Found acceptable parameter values are visualized in Figures S1, S2, and S3.

### Optimization and software

We used MATLAB R2019a (MathWorks, Natick, MA) and the IQM toolbox (IntiQuan GmbH, Basel, Switzerland) for modeling. The MATLAB-functions particleswarm and simulannealbnd were combined in the optimization runs for extensive searches of the space of parameters.

### Data processing

The number of repeats of each experiment was 2-6, and to save animals, the repeats did not come from new animals. Instead we used slices of lung from the same animal. We therefore believe that the true data uncertainty was underestimated. To correct for that, we used the maximal calculated SEM for each data series as a proxy for the actual SEM. We also had to use a correction factor in the model simulations to account for a scaling difference between the LPS-experiments and the Dexa-experiments (c.f. data in Figure 3B at dose 100 ng/ml LPS, and Figure 3C 100 ng/ml LPS at 24 hours in orange). This correction factor was included as an experimental model parameter and estimated in the range 1.5-1.9.

**Figure 3:**
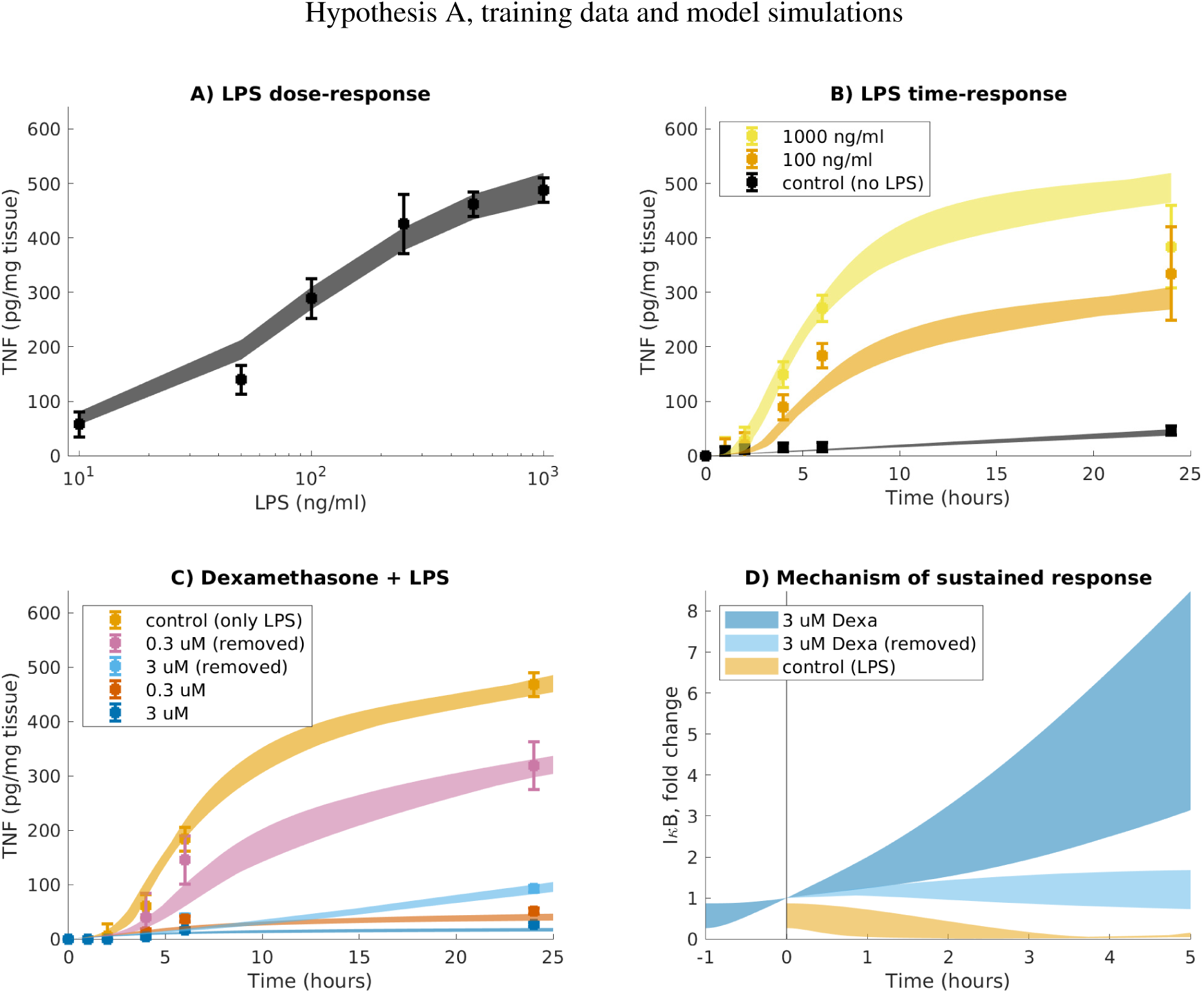
LPS with dexametasone pretreatments. Data and range of model simulations in agreement with data for Hypothesis A. Measured is TNF in response to different concentrations of LPS and/or Dexa. A) Different doses of LPS (10, 50, 100, 250, 500 and 1000 ng/ml) were used to trigger an inflammatory response, and the corresponding concentration of secreted TNF were measured after 24 hours. n = 2. B) 100 (yellow) and 1000 (orange) ng/ml LPS was added to trigger an inflammatory response and secreted TNF measured at several time points (0, 1, 2, 4, 6 and 24 hours). As control, no LPS was added (black). n = 6. C) Lung slices were pretreated with indicated concentrations of Dexa for 1 hour and next Dexa was either washed out (pink, sky blue), or not (red, blue). After that, 100 ng/ml LPS was added at time = 0. n = 3. D) The mechanism of sustained response is I*κ*B protein levels induced by Dexa in this hypothesis. I*κ*B binds to active NF-*κ*B and hinders transcription of TNF. Dots with error bars show data and standard errors of measurements, colored areas show the area of model simulations that are in agreement with data according to a *χ*^2^-test.

### Statistical analyses

For data comparison, we used One Way ANOVA with Tukey’s range test for multiple comparisons with a significance level of 0.05. To compute corresponding p-values, we used MATLAB-functions anova1 and multcompare. To reject models, we used the *χ*^2^-test [25] with a significance level of 0 .05. We used 37 degrees of freedom for training data (37 data points) leading to a threshold for rejection of *χ*^2^(0.05, 37) = 52, and 51 degrees of freedom for all data (51 data points) leading to a threshold for rejection of *χ*^2^(0.05, 51) = 69.

### Data and model availability

The experimental data as well as the complete code for data analysis and modeling are available at https://gitlab.liu.se/eliny61/macrophage-model.

### Animals

Male Wistar Han rats (Harlan, Horst, the Netherlands) weighing 225-300 gram were used in the present study. Animals were housed in an Association for Assessment and Accreditation of Laboratory Animal Care (AAALAC)-accredited animal facility in groups of six individuals at 18-22 degrees C under a 12-h light/dark cycle with access to water and chow ad libitum for at least 5 days prior to the experiments. The study was approved by the Animal Ethics Committee of Gothenburg (234–2011, 134–2013, and 137–2014).

### Preparation of lung slices

Lung slices were prepared from drug naive rat lungs as previously described [26]. Briefly, rat lungs were perfused via the pulmonary artery with saline and low melting point agarose solution and then inflated with low melting agarose solution via the trachea. The lobes were separated, and 500 *µ*m thick slices were made with DTH-Zero 1 Microslicer (Dosaka, Kyoto, Japan).

### Incubation of lung slices with LPS

Three lung slices were incubated in 15 mL buffer in 80-mm flat-bottomed glass dishes at 37 degrees C and 45 rpm in a Forma Orbital Shaker 420 (Thermo Fisher Scientific, Waltham, MA). Lung slices were incubated in different concentrations of LPS (10-1000 ng/ml, E.coli, 0.111:B4, Sigma, St Louis, MO, USA) during 24 hours in order to assess a dose-response relationship. Samples were collected from buffer (250 *µ*l) and protease inhibitor cocktail (4%, Thermo Scientific Pierce Protein Biology Products, Rockford, USA) was added. Slices were dried on filter paper, weighed, and 9 volumes (w/v) of PBS-buffer (140 mM NaCl, 10 mM phosphate buffer, 3 mM KCl, Calbiochem, U.S. and Canada) were added. The samples were supplemented with protease inhibitor cocktail (4%) prior to homogenization with an ultrasonic probe (Sonifier 250; Branson Ultrasonics, Danbury, CT). Lung slices were also incubated with 10, 100 and 1000 ng/ml LPS and sampling from the supernatants was performed at different time points to evaluate time-dependent release of TNF.

### Incubation of lung slices with LPS and Dexa

Lung slices were pre-loaded with Dexa (Sigma) in two different concentrations (0.3 and 3 *µ*M) one hour prior LPS-stimulation. Half of the setup received Dexa as a one hour pre-treatment while the remaining were given a continuous supply of Dexa throughout the LPS provocation in addition to the one hour pre-treatment. Lung slices that received a short-term exposure of Dexa underwent one buffer exchange in order to remove residual Dexa from the surface of the slices. Samples from buffer were collected (250 *µ*l) and supplemented with protease inhibitor cocktail (4%) at time points 0, 2, 4, 6 and 24 hours.

Lung slices were also pre-treated with Dexa (3 *µ*M) during one hour at different time points followed by three buffer changes. After buffer exchange, the slices were kept in lung slice buffer in different time intervals before stimulation with LPS (100 ng/ml). Samples were collected from supernatants (250 *µ*l) and added with protease inhibitor cocktail (4%) at 0, 1, 2, 4 and 6 hours after LPS stimulation.

### TNF quantification

Enzyme-linked immunosorbent assay (ELISA DuoSet®, R&D Systems, Abingdon, UK) was used to quantify the levels of TNF in supernatants and lung slice homogenate according to manufacturer’s instructions. Levels of quantification (LOQ) were 62.5-4000 pg/ml.

## Results

### Data collection

To unravel a potential sustained anti-inflammatory effect of Dexa in lung slices, we combined experimental work and mathematical modeling. We used sliced lung tissue from rats in all experiments. First, we collected data for different concentrations of LPS after 24 hours of stimulation (Figure 3A, dots with error bars). LPS induced activation of inflammation, as measured by TNF supernatant levels. Maximal activation was achieved at 500-1000 ng/ml of LPS. Intracellular levels of TNF did not show a dose-response relationship with LPS (data not shown), and we therefore focused subsequent experiments on extracellular levels of TNF. We performed a total protein determination in the lung tissue slices, and the protein content in the samples correlated to the weight of the samples (data not shown) which allowed data to be weighted with the mass of the tissue slices. Second, we collected time-resolved data for control + LPS induced increase in TNF for 100 and 1000 ng/ml LPS (Figure 3B, dots with error bars). LPS induced a detectable response at TNF after 2 hours, a linear increase between 2-6 hours, and a saturation somewhere before 24 h. Third, we collected data for Dexa-induced suppression of TNF secretion (Figure 3C, dots with error bars). Dexa was added for 1 hour before the addition of LPS. At the time of LPS addition, Dexa was either kept in the solution or washed away. In these experiments with Dexa, we used a sub-maximal concentration of LPS, 100 ng/ml, to be able to study potential suppression Hypothesis A, training data and model simulations of the inflammatory response. Indeed, Dexa suppressed the LPS-induced secretion of TNF, even when removed from the solution before addition of LPS (Figure 3C, pink and sky blue). A high dose of Dexa (3 *µ*M), that was removed from the solution before addition of LPS, suppressed the TNF release to similar levels as a low dose of Dexa (0.3 *µ*M) that was kept in the solution (cf. sky blue and red in Figure 3C, mean values are 91 and 51 pg/mg tissue). In summary, our data shows a sustained anti-inflammatory effect of a high dose of Dexa that has been washed away from the lung slices.

### Hypothesis testing

To go from data to mechanistic insights, we developed mathematical models based on these data, for both Hypothesis A: “New synthesis of I*κ*B*α*” and for Hypothesis B: “Direct inactivation of NF-*κ*B” (Figure 1). The two different models were based on known signaling mechanisms and existing knowledge of LPS-induced activation of inflammation and Dexa-induced reduction of inflammation (e.g. [11, 13]), and differed only in the implementation of the effect of Dexa. The models were kept small, including only key mechanisms (Figure 2), to reduce the number of model parameters to estimate from the limited data. Both Hypothesis A (Figures 3A-C) and Hypothesis B (Figures 4A-C) showed a good agreement with estimation data (c.f. dots and lines in the same colors in the figures). We collected sets of acceptable parameters for both hypotheses by running the parameter estimation procedure multiple times and saving parameters that gave a good agreement between model simulations and data according to the statistical *χ*^2^-test [25] with level of significance 0.05 and 37 degrees of freedom (37 data points). In this way we could get an approximation for the uncertainty of predictions by the model. These uncertainties are displayed as colored areas in the figures. The found parameter values are visualized in Figures S1 and S2.

**Figure 4:**
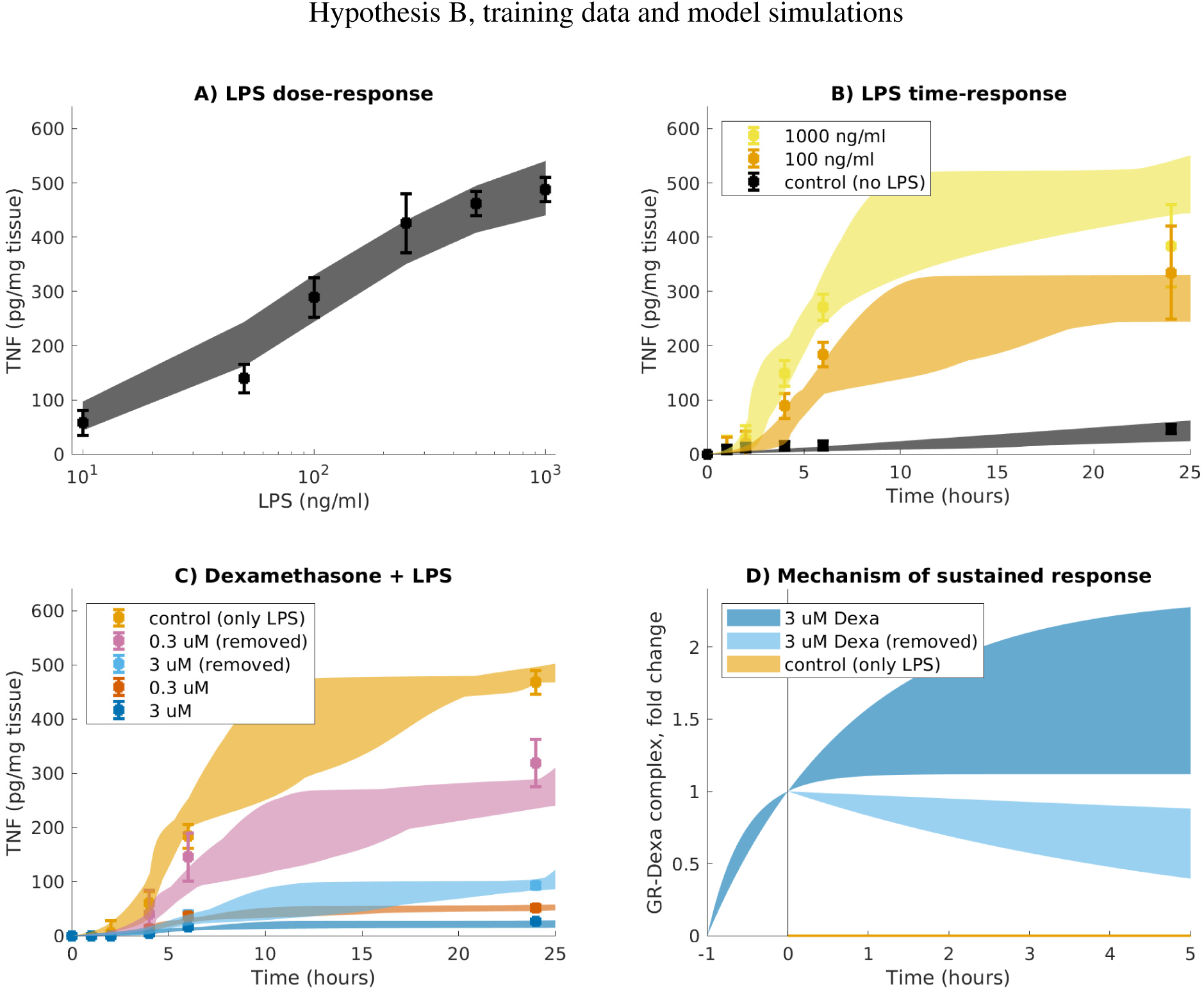
Data and range of model simulations in agreement with data for Hypothesis B. Measured is TNF in response to different concentrations of LPS and/or Dexa. A) Different doses of LPS (10, 50, 100, 250, 500 and 1000 ng/ml) were used to trigger an inflammatory response, and the corresponding concentration of secreted TNF were measured after 24 hours. n = 2. B) 100 (yellow) and 1000 (orange) ng/ml LPS was added to trigger an inflammatory response and secreted TNF measured at several time points (0, 1, 2, 4, 6 and 24 hours). As control, no LPS was added (black). n = 6. C) Lung slices were pretreated with indicated concentrations of Dexa for 1 hour and next Dexa was either washed out (pink, sky blue), or not (red, blue). After that, LPS was added at time = 0. n = 3. D) The mechanism of sustained response is at the level of the GR-Dexa complex in this hypothesis. The slow release of Dexa from GR allow for continuous inhibition of NF-*κ*B transcriptional activity. Dots with error bars show data and standard errors of measurements, colored areas show the area of model simulations that are in agreement with data according to a *χ*^2^-test.

### Mechanisms of sustained response

We used these areas to study the mechanism of the sustained response to Dexa of the two hypotheses. For Hypothesis A, the level of I*κ*B protein, as induced by Dexa, is responsible for the sustained response (Figure 3D). When Dexa is removed, I*κ*B remains (cf. sky blue area with the control in orange that goes down to 0 in Figure 3D). I*κ*B binds to active NF-*κ*B and hinders transcription of TNF. For hypothesis B, the mechanism of sustained response is at the level of the GR-Dexa (Figure 4D). When Dexa is removed, the GR-Dexa remains active for a long time (see the sky blue area in Figure 4D). The slow release of Dexa from GR allow for continuous inhibition of NF-*κ*B transcriptional activity.

### Test with new data

Both Hypothesis A and B were evaluated further to see their predictive abilities. To do so, we simulated the response to 100 ng/ml LPS and 3 *µ*M Dexa. This time, we used increasing lengths of the wash between Dexa and LPS addition, to see how long the response to Dexa was predicted to be sustained. Within a 30 hour time period, Hypothesis A showed a similar, totally sustained, response regardless of the length of the wash (Figure 5A). Hypothesis B, on the other hand, showed a more differential behavior, where the length of the wash affected the response (Figure 6A). Note that not all acceptable parameters in Hypothesis B are predicting a sustained response. Model simulations with parameters that give rise to a sustained response over time according are indicated in Figure 6A with darker areas.

**Figure 5:**
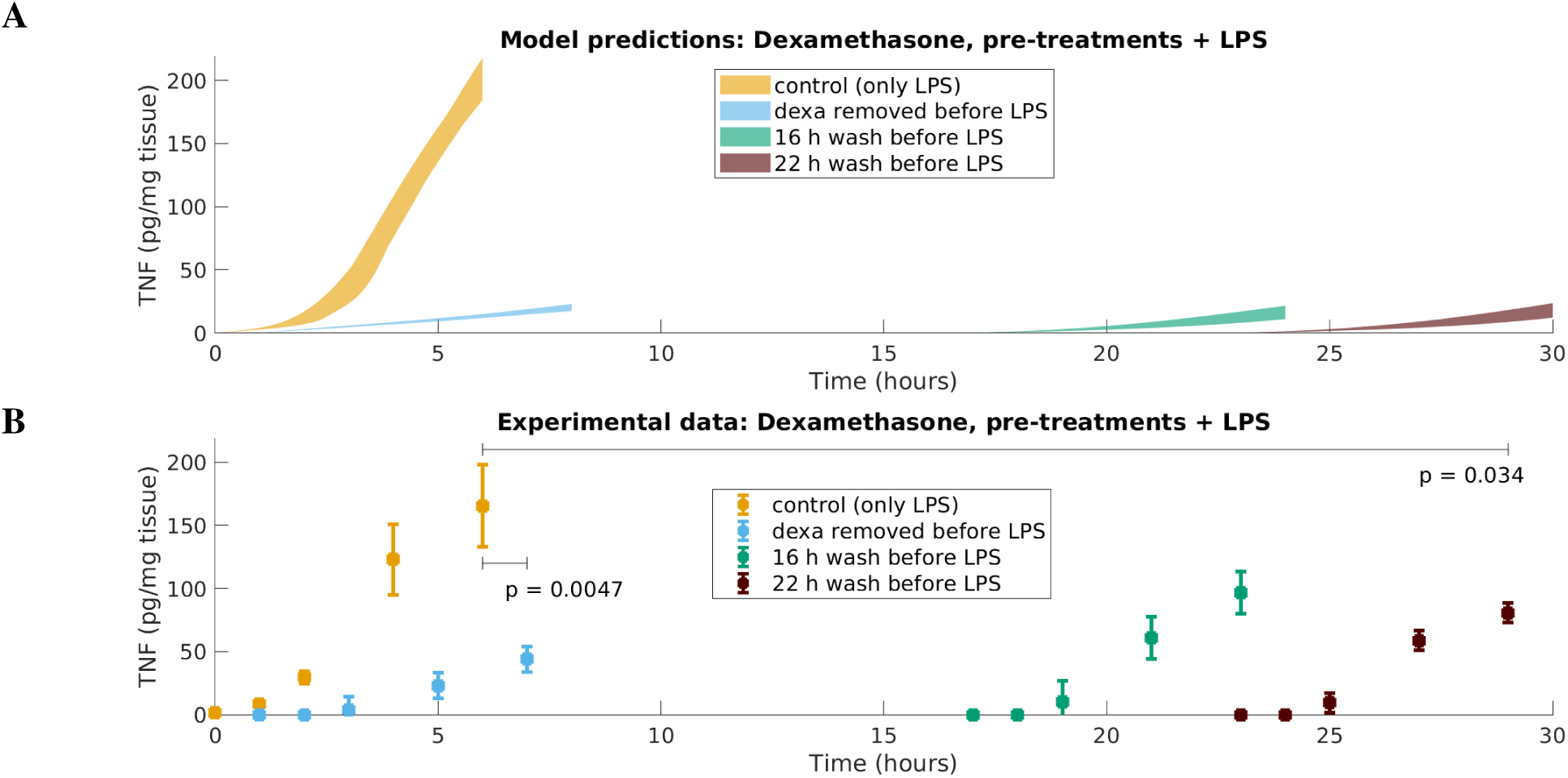
Rejection of Hypothesis A: “New synthesis of I*κ*B*α*”. A) The range of model simulations for Hypothesis A in agreement with data in Figure 3 shows a predicted response to Dexa of TNF that is totally sustained even after 22 hours of wash. B) Measured TNF in response to different concentrations of LPS and/or Dexa as indicated. Lung slices were pretreated with Dexa for 1 hour before washout during indicated times. At time = 0, 100 ng/ml LPS is added. n = 3. Dots with error bars show data and standard errors of measurements. The model simulations in A) is not in agreement with these data and must be rejected. Colored areas in A) show the area of model simulations that are in agreement with data in Figure 3 according to a *χ*^2^-test.

**Figure 6:**
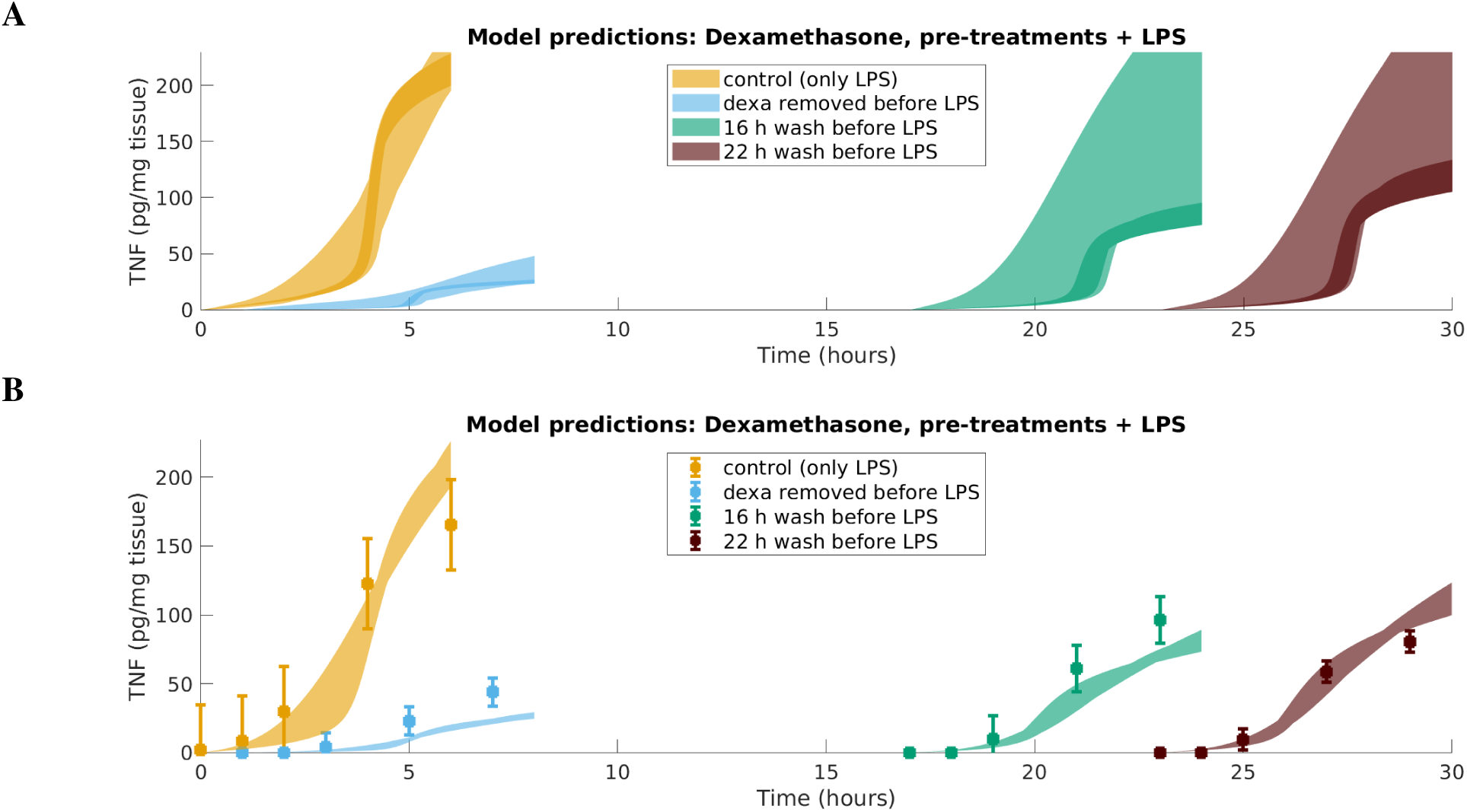
In agreement with new data, Hypothesis B: “New synthesis of I*κ*B*α*”. A) The range of model simulations for Hypothesis B in agreement with data in Figure 4 shows a predicted response to Dexa of TNF that cannot be rejected based on corresponding data in B). Highlighted are selected model simulations that show a sustained response. B) Model simulations in agreement with all data, i.e. the model has been trained to fit with both the data here and the data in Figure S4. Dots with error bars show data and standard errors of measurements.

We performed the corresponding measurements experimentally to test these predictions (Figure 5B, see Method for experimental details), and corresponding data showed a clear time-dependency in the sustainability of the response – the longer the wash, the less sustained the response (Figure 5B). When Dexa was removed just before the addition of LPS, the TNF release was inhibited to a large degree (Figure 5B, sky blue). When Dexa was removed after 16 or 22 hours of wash, there was still an inhibition of TNF release, however not as strong (Figure 5B, green and brown) as compared to control (Figure 5B, orange) Therefore, Hypothesis A had to be rejected. We also ran the optimization again, including both training and testing data, to assure that there were no parameter set for hypothesis A that we had missed. Despite many searches, we could not find any parameter Hypothesis B, training data and model simulations set in agreement with all data for Hypothesis A.

Hypothesis B, on the other hand, showed a more differential behavior, where the length of the wash affected the response (Figure 6A). Note that not all acceptable parameters in Hypothesis B are predicting a sustained response. The parameters in agreement with both the training data and the test data, i.e. the parameters that gave a sustained response over time according to data, are indicated in Figure 6A with darker areas. For further analysis, we searched for more parameters in agreement with both training and test data for Hypothesis B. A new threshold for the *χ*^2^-test was calculated with 51 degrees of freedom, since all data contains 51 data points. This new set of parameters (Figure S3), in agreement with all data (Figure 6B and S4), is used to simulate treatment schemes below.

### Treatment schemes

To display the potential of the developed model, we next used the accepted hypothesis, together with the found parameters in agreement with all data, to simulate hypothetical treatment schemes. We assumed patients with some inflammatory disease, displayed by high levels of TNF at the site of inflammation. We simulated the disease by adding a constant infusion delivering 1 ng/mL LPS at steady state to the model and ran the simulation to steady state. To account for the different situation in plasma compared to the situation in vitro under which the model was developed, we added two parameters to the model. The first added parameter accounted for the half life of Dexa in plasma. We assumed this half life to be 2 hours, in accordance with Queckenberg et al. [27]. The second added parameter accounted for the rate of elimination of TNF, which we assumed to be per hour, a value taken from Held et al. [28]. We simulated four different treatment schemes with Dexa in such patients (Figure 7), with the same total daily dose of Dexa (corresponding to 0.3 *µ*M elevation of Dexa concentration in plasma): One treatment per day (0.3 *µ*M Dexa), Two treatments per day (0.15 + 0.15 *µ*M Dexa), Three treatments per day (0.1 + 0.1 + 0.1 *µ*M Dexa), and continuous 24 treatments per day (0.3/24 *µ*M Dexa per occasion). See Figure 7, right, for the dosing scheme. The model simulations show that the sustained response to Dexa, which the model captures, allow for an optimal treatment scheme already at one or two treatments per day (Figure 7, left). We calculated the area under the curve (AUC) with uncertainties for the TNF levels during the second day of treatment. We selected the area between then model simulations and the maximal response (lower horizontal line in Figure 7A). The second day of treatment was chosen to remove the effect of the rate of the decrease from a rather high level before treatment (Figure 7, left). One treatment per day gives an interval of AUC = (0.71, 1.21). Two treatments per day gives a small improvement, AUC = (0.52, 0.99). Three treatments (or more) per day gives no further improvement, AUC = (0.50, 0.97). The explanation to the sustained response is the slow release of Dexa from the Dexa-GR complex (Figure 7, middle), as we know from previous analyses cf. Figure 4D.

**Figure 7:**
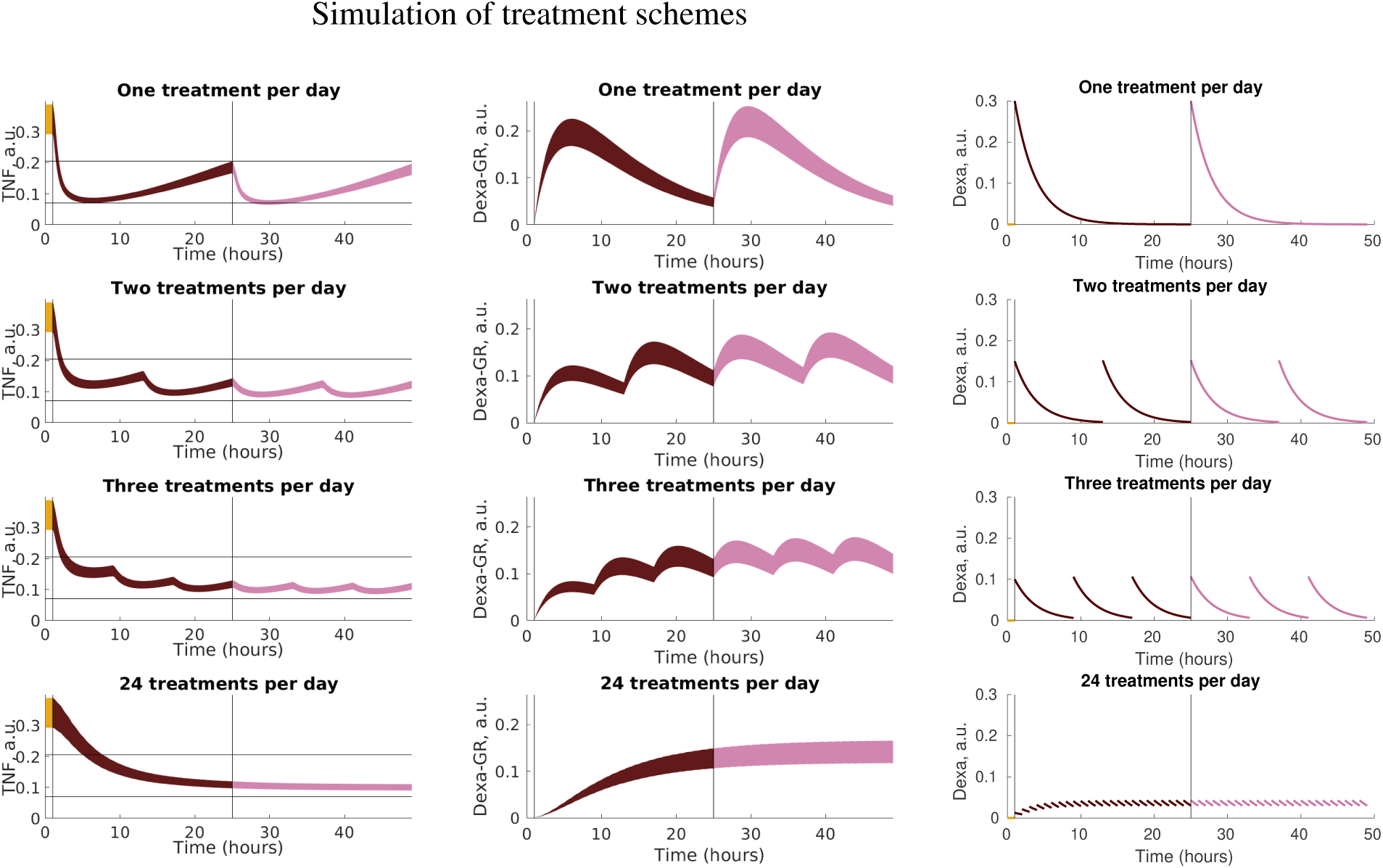
Simulation of treatment schemes under the assumption that the half life of Dexa is 2 hours and that TNF is eliminated with the rate 5.65 per hour [28]. The same total dose of drug is subdivided into 1-24 treatments. Left: TNF levels in plasma. With one treatment per day the effect of Dexa is already rather sustained, AUC = (0.71, 1.21). Two treatments per day gives a small improvement, AUC = (0.52, 0.99). Three or more treatments per day gives no further substantial improvement: AUC = (0.50, 0.97) and AUC = (0.52, 1.00). AUC is calculated the second day of treatment. The horizontal black lines indicate the maximal and minimal TNF levels for the one treatment per day scheme. The vertical black lines indicate start of a new day. Middle and Right: The formation of Dexa-GR complex and the dynamics of Dexa under the different treatment schemes.

## Discussion

We have analyzed mechanisms of a sustained anti-inflammatory response in alveolar macrophages. To do so, we have combined experimental data and a mechanistic modeling approach. Our main findings is that i) in alveolar macrophages, there is a sustained anti-inflammatory response to Dexa that can be explained by a slow release of Dexa from the Dexa-GR complex; ii) one of the main hypotheses for the intracellular effect of Dexa, Hypothesis A: “New synthesis of I*κ*B*α*”, cannot explain our data, instead the alternative hypothesis, Hypothesis B: “Direct inactivation of NF-*κ*B”, can explain all our data; iii) our final model can be used to simulate hypothetical treatment scenarios to see that a once-daily dose of Dexa would be enough to obtain a beneficial anti-inflammatory response over 24 hours.

Our conclusions in ii) are in line with reports detailing how Dexa inhibit NF-*κ*B signaling, not primarily through affecting I*κ*B*α* (Hypothesis A) but rather by interacting with the glucocorticoid receptor (GR) and affecting NF-*κ*B in the nucleus. Direct interaction between GR and p65 has been suggested through immunoprecipitation experiments [13], but the suppression of NF-*κ*B activity is likely mediated through downstream events such as increased export from the nucleus or site-specific phosphorylation of NF-*κ*B subunits [29]. It has been shown previously that Dexa, after binding to GR, interacts with NF-*κ*B and promotes enhanced export of the p65 subunit from the nucleus, peaking at 20 min [30]. The sustained response observed in the current study may include GR-induced expression of additional proteins such as Glucocorticoid-induced leucine zipper (GILZ), that both bind and inhibit p65 as well and further stimulates increased export of p65 out of the nucleus [31, 32].

We have used a mechanistic modeling approach, in contrast to an empirical PK/PD modeling approach which is more often used in pharmacological applications. The benefit of a mechanistic modeling approach is that we can not only predict left out data, which is possible with both approaches, but also study the mechanisms behind observed phenomenon. In this study, we used mechanistic modeling first to distinguish between two biological hypotheses for the intracellular effect of Dexa. Such hypothesis testing allow for conclusions in the form of rejections, and predictions with uncertainties that can be tested experimentally [25]. We find a model prediction that distinguish between the biological hypotheses, and corresponding measurement shows that one of the hypotheses cannot explain these data. This is an excellent example of how mechanistic modeling can be used in experimental design, to assure that new esperiments give rise to new conclusions about the biological system under study. Second, we used mechanistic modeling to gain new insights into the mechanisms of the sustained response. For the two hypotheses, we found different mechanisms that were responsible for the sustained response: an increase in I*κ*B protein in response to Dexa (Figure 3D) and a slow release of Dexa from the receptor (Figure 4D). Finally, we combined our final mechanistic model with a few key PK parameters to simulate a hypothetical scenario of treatment schemes. Such a combined model is commonly referred to as a systems pharmacology model and make use of the strengths of both modeling approaches. The included mechanistic details gives insights also in the simulation of treatment scenarios, e.g. the mechanism of the sustained response herein (Figure 7B).

We assume that we have a complete wash of Dexa from the lung slices. Theoretical calculations involving known characteristics of Dexa shows that 0-2 % of Dexa might bind to the tissue and thus not be washed away. To account for such (small) effects, we have preformed parameter estimation with both hypotheses with corresponding assumptions, i.e. that we do not have a complete wash. The conclusions herein do not change if we change the assumption to allow for not complete washout (not shown).

We provide a model of the sustained effect of an anti-inflammatory drug in alveolar macrophages. The mechanistic modeling approach allow for incremental model extensions whenever more data becomes available. Interesting extensions for the model would be to add other compounds with known or unknown mechanisms of actions, more intracellular signaling details, as well as connecting the macrophages model to models for other players in the immune system. Our work opens up for more systematic searches for drug candidates with sustained anti-inflammatory responses, which could lead to smaller variations in drug concentrations, and thus potentially to drugs with fewer side effects.

## List of abbreviations

TNF: Tumor necrosis factor;
TLR: toll like receptor;
NF-*κ*B: nuclear factor *κ*-light-chain-enhancer of activated B cells;
I*κ*B*α*: inhibitor of NF-*κ*B;
IKK: I*κ*B kinase;
IRAK: Interleukin-1 receptor-associated kinase;
TRAF: TNF receptor associated factor;
Dexa: Dexamethasone;
GR: glucocorticoid receptor;
LPS: lipopolysaccharides;
PK/PD: pharmacokinetic/pharmacodynamic.

## Supplemental Material

**Figure S1:**
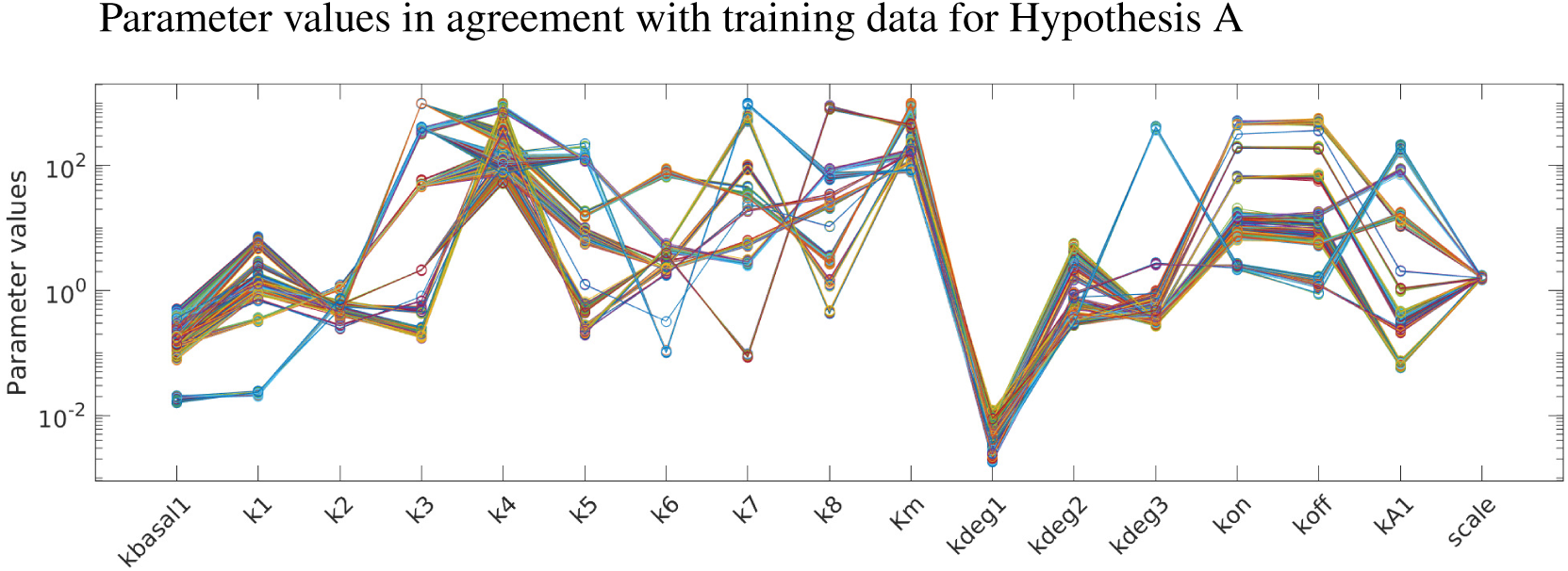
A representation of the acceptable parameter values for Hypothesis A that give rise to the simulations in agreement with training data in Figure 3.

**Figure S2:**
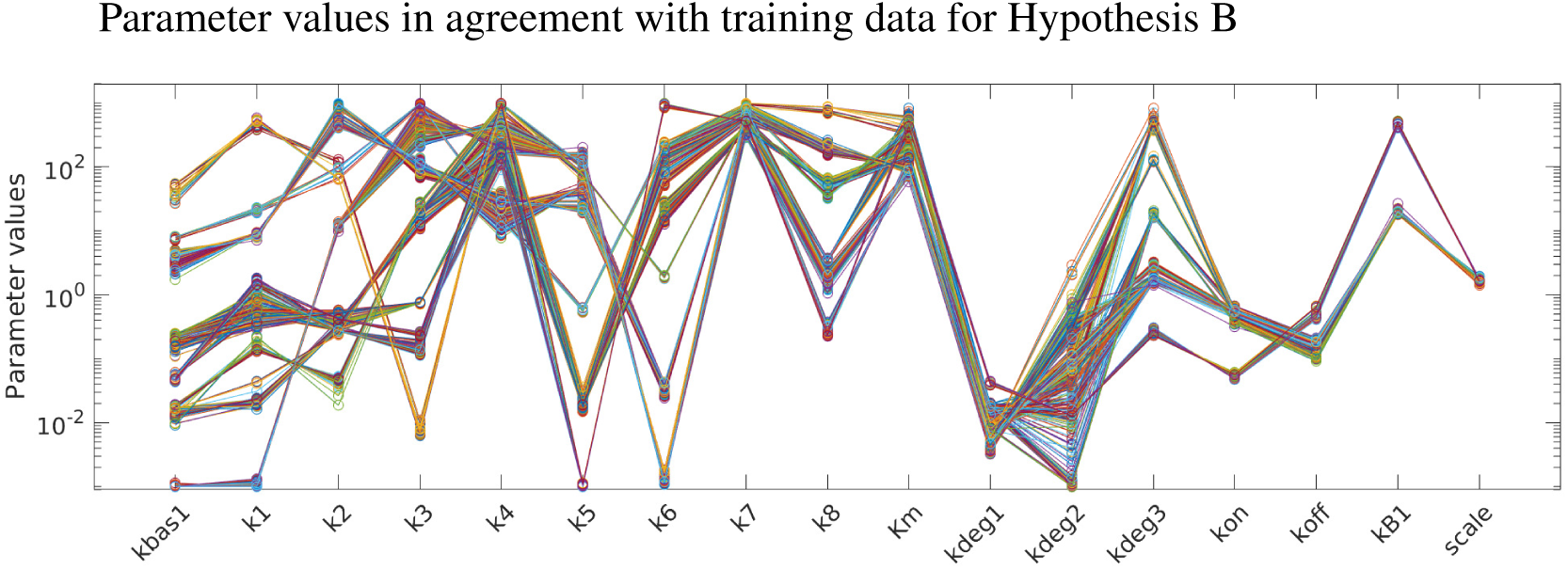
A representation of the acceptable parameter values for Hypothesis B that give rise to the simulations in agreement with training data in Figure 4.

**Figure S3:**
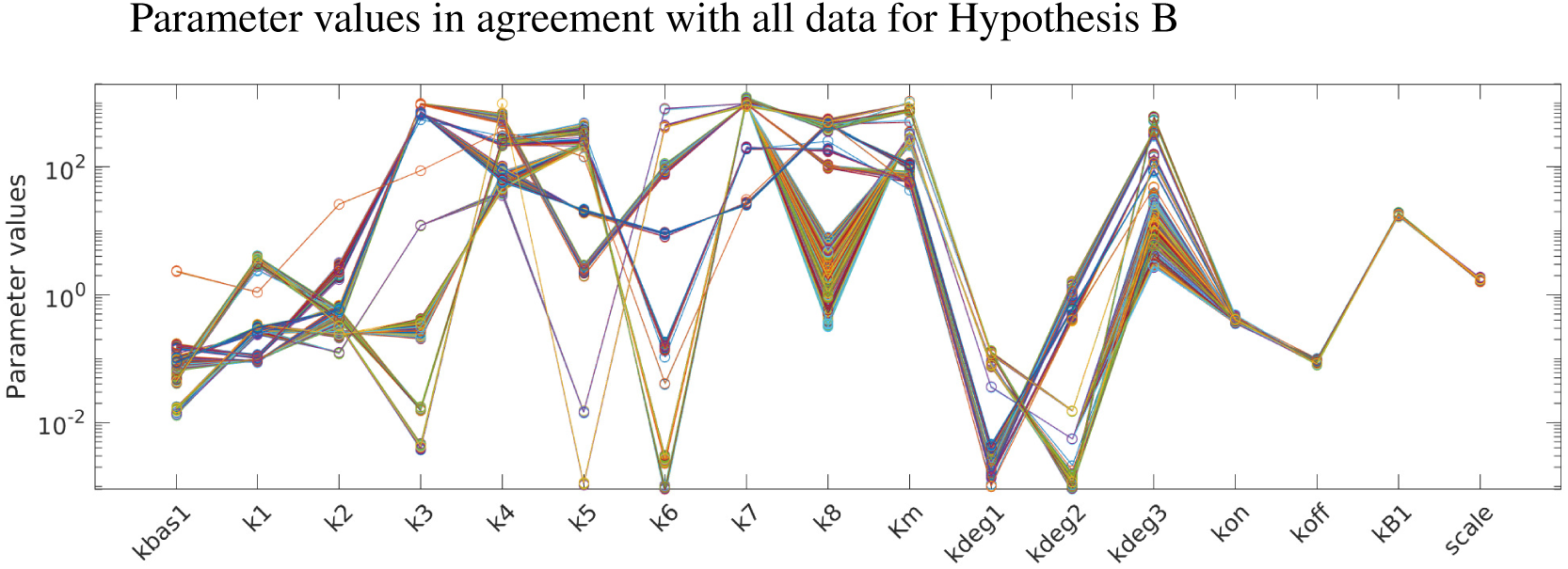
A representation of the acceptable parameter values for Hypothesis B that give rise to the simulations in agreement with all data in Figures S4 and 6B

**Figure S4:**
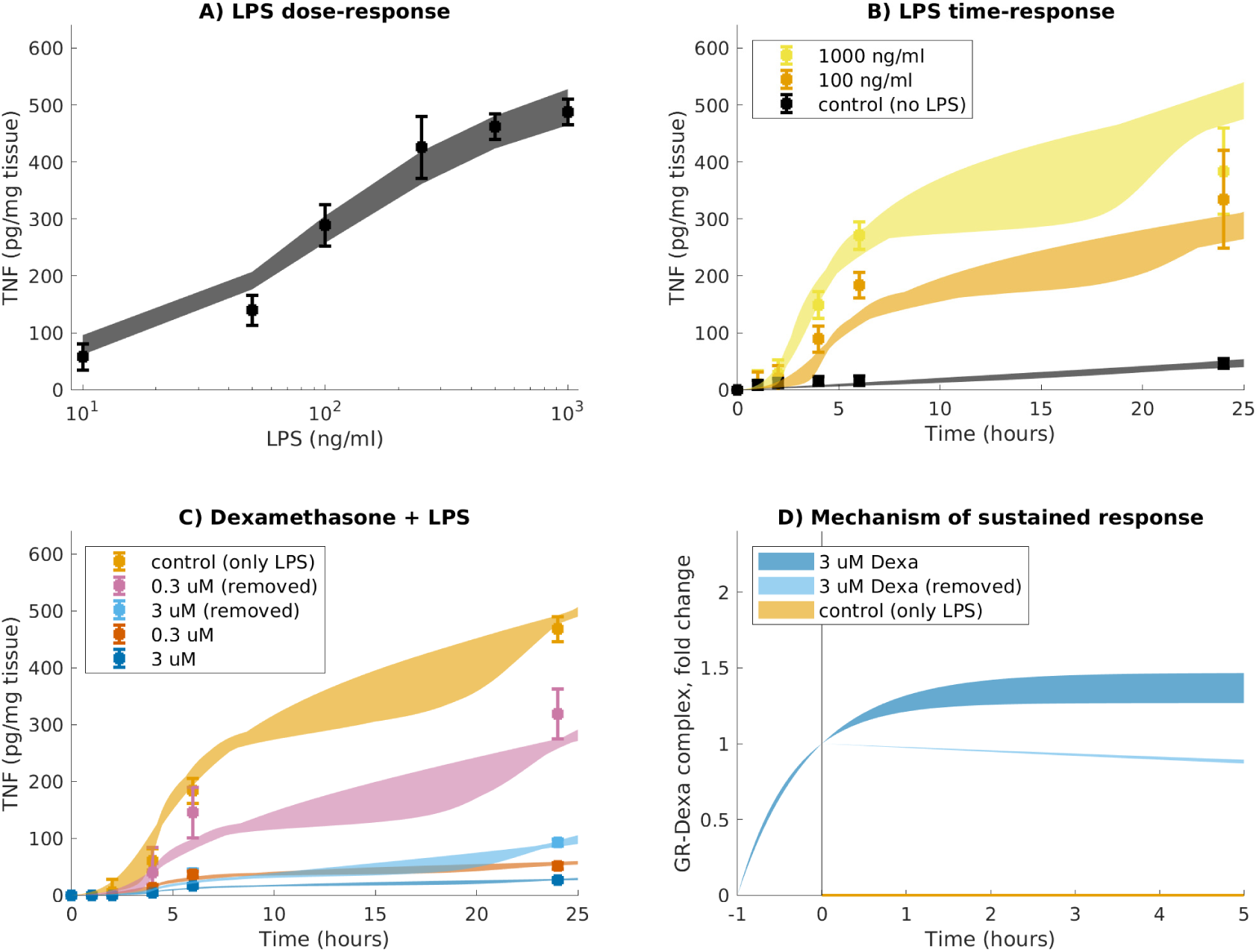
Data and range of model simulations in agreement with all data for Hypothesis B. Measured is TNF in response to different concentrations of LPS and/or Dexa. A) Different doses of LPS (10, 50, 100, 250, 500 and 1000 ng/ml) were used to trigger an inflammatory response, and the corresponding concentration of secreted TNF were measured after 24 hours. n = 2. B) 100 (yellow) and 1000 (orange) ng/ml LPS was added to trigger an inflammatory response and secreted TNF measured at several time points (0, 1, 2, 4, 6 and 24 hours). As control, no LPS was added (black). n = 6. C) Lung slices were pretreated with indicated concentrations of Dexa for 1 hour and next Dexa was either washed out (pink, sky blue), or not (red, blue). After that, LPS was added at time = 0. n = 3. D) The mechanism of sustained response is at the level of the GR-Dexa complex in this hypothesis. The slow release of Dexa from GR allow for continuous inhibition of NF-*κ*B transcriptional activity. Dots with error bars show data and standard errors of measurements, colored areas show the area of model simulations that are in agreement with data according to a *χ*^2^-test.

